# A novel rubi-like virus in the Pacific electric ray (*Tetronarce californica*) reveals the complex evolutionary history of the *Matonaviridae*

**DOI:** 10.1101/2021.02.08.430315

**Authors:** Rebecca M. Grimwood, Edward C. Holmes, Jemma L. Geoghegan

## Abstract

*Rubella virus* (RuV) is the causative agent of rubella (“German measles”) and remains a global health concern. Until recently, RuV was the only known member of the genus *Rubivirus* and the only virus species classified within the *Matonaviridae* family of positive-sense RNA viruses. Other matonaviruses, including two new rubella-like viruses, *Rustrela virus* and *Ruhugu virus*, have been identified in several mammalian species, along with more divergent viruses in fish and reptiles. To screen for the presence of additional novel rubella-like viruses we mined published transcriptome data using genome sequences from *Rubella, Rustrela*, and *Ruhugu viruses* as baits. From this, we identified a novel rubella-like virus in a transcriptome of *Tetronarce californica* (Pacific electric ray) that is more closely related to mammalian *Rustrela virus* than to the divergent fish matonavirus and indicative of a complex pattern of cross-species virus transmission. Analysis of host reads confirmed that the sample analysed was indeed from a Pacific electric ray, and two other viruses identified in this animal, from the *Arenaviridae* and *Reoviridae*, grouped with other fish viruses. These findings indicate that the evolutionary history of the *Matonaviridae* is more complex than previously thought and highlights the vast number of viruses still to be discovered.

## 1. Introduction

*Rubella virus* (RuV) (*Matonaviridae: Rubivirus*) is a single-stranded positive-sense RNA virus [1]. It is best known as the causative agent of rubella, sometimes known as “German measles” -a relatively mild measles-like illness [2]. More severely, RuV is also teratogenic and can result in complications such as miscarriage or congenital rubella syndrome (CRS) if contracted during pregnancy [3, 4]. Despite the availability of effective vaccines, including the combination measles, mumps, and rubella (MMR) vaccine [5], they are deployed in only around half of the world’s countries [6]. RuV therefore remains a significant global health concern, particularly in developing countries where childhood infection rates are high and vaccination efforts are low or non-existent [7].

The disease rubella was first described in 1814 [8] and until recently RuV was the only known species within the genus *Rubivirus*, itself the only genus in the family *Matonaviridae* [9]. This picture has changed with the discovery of other rubella-like viruses through metagenomic technologies. The first of these, *Guangdong Chinese water snake rubivirus*, was discovered via a large virological survey of vertebrates in China and is highly divergent from RuV, sharing around 34% nucleotide similarity [10]. *Tiger flathead matonavirus* was identified through a meta-transcriptomic exploration of Australian marine fish [11], which, along with *Guangdong Chinese water snake rubivirus*, forms a sister clade to RuV. Even more recently, two novel rubella-like virus species were described: *Rustrela virus*, identified in several mammalian species close by and in a zoo in Germany, and *Ruhugu virus* from bats in Uganda [12]. These viruses represent the first non-human mammalian viruses within this family. Such findings begin to shed light on the deeper evolutionary history of this previously elusive viral family.

The introduction and popularisation of metagenomic methods to identify novel viruses within tissue or environmental samples has enabled their rapid discovery and provided important insights into viral diversity and evolution [13, 14]. Here, we performed data-mining of metagenomic data to determine if related matonviruses were present in other vertebrate taxa, screening the National Center for Biotechnology Information (NCBI) Transcriptome Shotgun Assembly (TSA) database against the genome sequences of *Rubella, Ruhugu*, and *Rustrela viruses*.

## 2. Materials and Methods

### 2.1 TSA mining for novel matonaviruses

To identify the presence of novel rubella-like viruses in vertebrates, translated genome sequences of *Rustrela virus* (MN552442, MT274274, and MT274275), *Ruhugu virus* (MN547623), and RuV strain 1A (KU958641) were screened against transcriptome assemblies available in NCBI’s TSA database (https://www.ncbi.nlm.nih.gov/genbank/tsa/) using the translated Basic Local Alignment Search Tool (tBLASTn) algorithm [15]. Searches were restricted to vertebrates (taxonomic identifier: 7742), excluding *Homo sapiens* (taxonomic identifier: 9606) and utilised the BLOSUM45 matrix to increase the chance of finding highly divergent viruses. Putative viral sequences were then queried using the DIAMOND BLASTx (v.2.02.2) [16] algorithm against the non-redundant (nr) protein database for confirmation.

### 2.2 Pacific electric ray transcriptome assembly and annotation

To recover further fragments of the novel matonavirus genome and screen for other viruses in the Pacific electric ray transcriptome (see Results), raw sequencing reads were downloaded from the NCBI Short Read Archive (SRA) (Bioproject: PRJNA322346). These reads were first quality trimmed and then assembled *de novo* using Trinity RNA-Seq (v2.11) [17]. Assembled contigs were annotated based on similarity searches against the NCBI nucleotide (nt) database using the BLASTn algorithm and the non-redundant protein (nr) database using DIAMOND BLASTx (v.2.02.2) [16]. Novel viral sequences were collated by searching the annotated contigs and these potential viruses were confirmed with additional BLASTx searches against nr and nt databases.

### 2.3 Abundance estimations

Virus abundance, expressed as the standardised total number of raw viral sequencing reads that comprise a given contig, was calculated using the ‘align and estimate’ module available for Trinity RNA-seq with RNA-seq by Expectation-Maximization (RSEM) [18] as the abundance estimation method and Bowtie 2 [19] as the alignment method.

### 2.4 Phylogenetic analysis

A range of virus genomes representative of each virus family and sub-family under consideration, and for a range of host species, were retrieved from GenBank for phylogenetic analysis. The amino acid sequences of previously described viruses were aligned with those generated here with Multiple Alignment using Fast Fourier Transform (MAFFT) (v.7.4) [20] employing the E-INS-i algorithm. All sequence alignments were visualised in Geneious Prime (v2020.2.4) [21]. Ambiguously aligned regions were removed using trimAl (v.1.2) [22] employing the ‘gappyout’ flag. The maximum likelihood approach available in IQ-TREE (v.1.6.12) [23] was used to generate phylogenetic trees for each family. ModelFinder [24] was used to determine the best-fit model of amino acid substitution for each tree and 1000 ultrafast bootstrap replicates [25] were performed to assess nodal support. Phylogenetic trees were annotated with FigTree (v.1.4.4) [26].

### 2.5 Virus nomenclature

The new *T. californica* viruses described were named with ‘*Tetronarce*’ preceding their respective family name to signify their electric ray host.

### 2.6 Data availability

The *Rustrela* and *Ruhugu virus* sequences used for data mining in the project are available under Bioproject PRJNA576343. The raw *T. californica* sequence reads are available at Bioproject PRJNA322346. All other publicly available sequences used in analyses can be accessed via their corresponding accession codes (see relevant figures). Consensus sequences of the new viruses identified in this study will be available on GenBank.

## 3. Results

### 3.1 Identification of a novel Matonaviridae in Pacific electric ray

We identified a complete structural polyprotein and several fragments of the RNA-dependent RNA polymerase (RdRp) from a divergent rubi-like virus that we tentatively named *Tetronarce matonavirus*. Screening of *Rustrela, Ruhugu*, and *Rubella 1A virus* sequences against the TSA database revealed matches to a contig of 3006 nucleotides (1001 amino acids) from a transcriptome of the electric organ of a female Pacific electric ray (*Tetronarce californica*) [27], ranging in sequence identity from 37.6 – 44.9%. Querying this contig against the non-redundant protein database using the BLASTx algorithm confirmed that it had ∼38% sequence identity to the structural protein of *Rubella virus* (accession: AAY34244.1, query coverage: 87%, e-value 0.0, see Table 1).

**Table 1.**
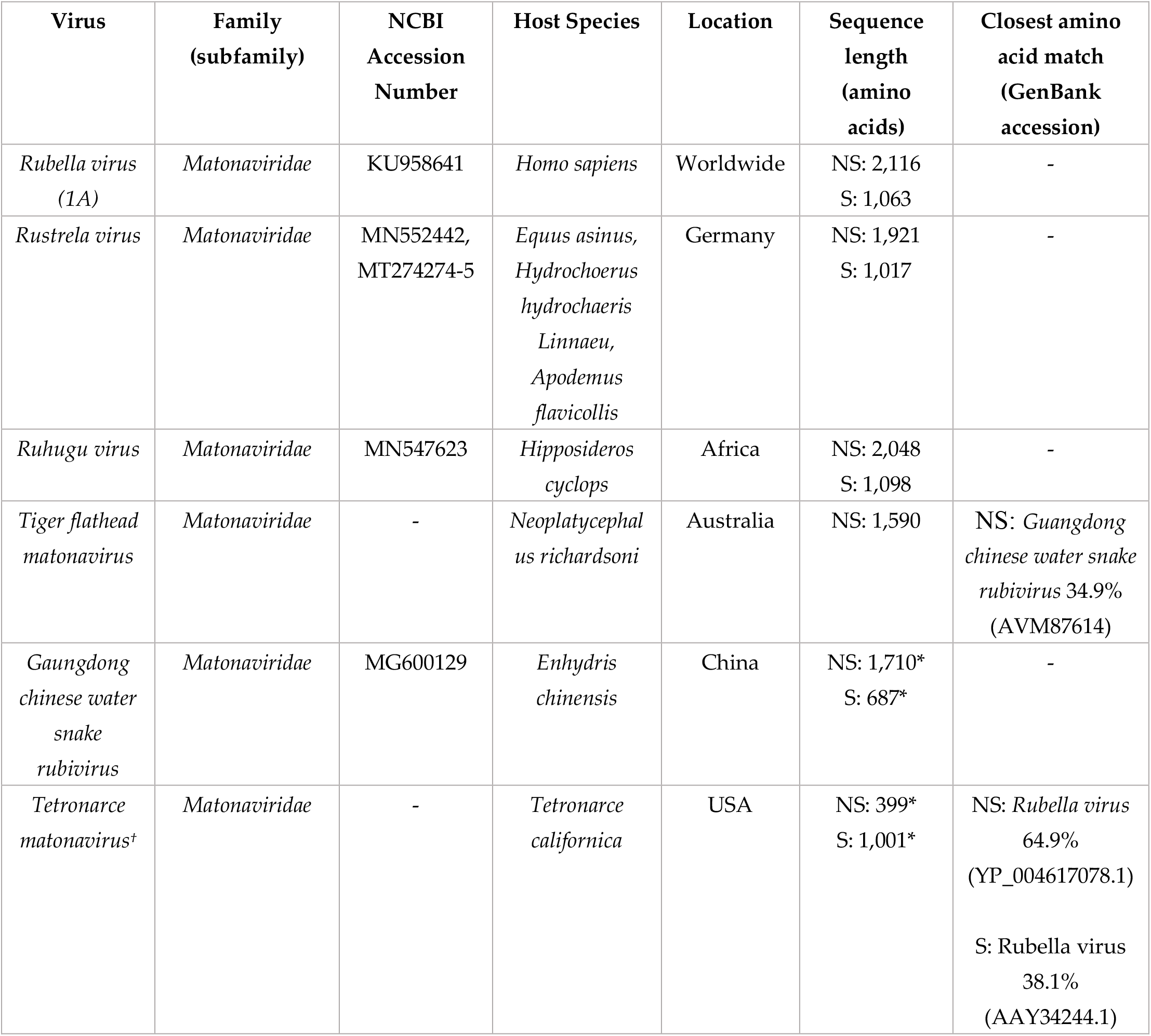
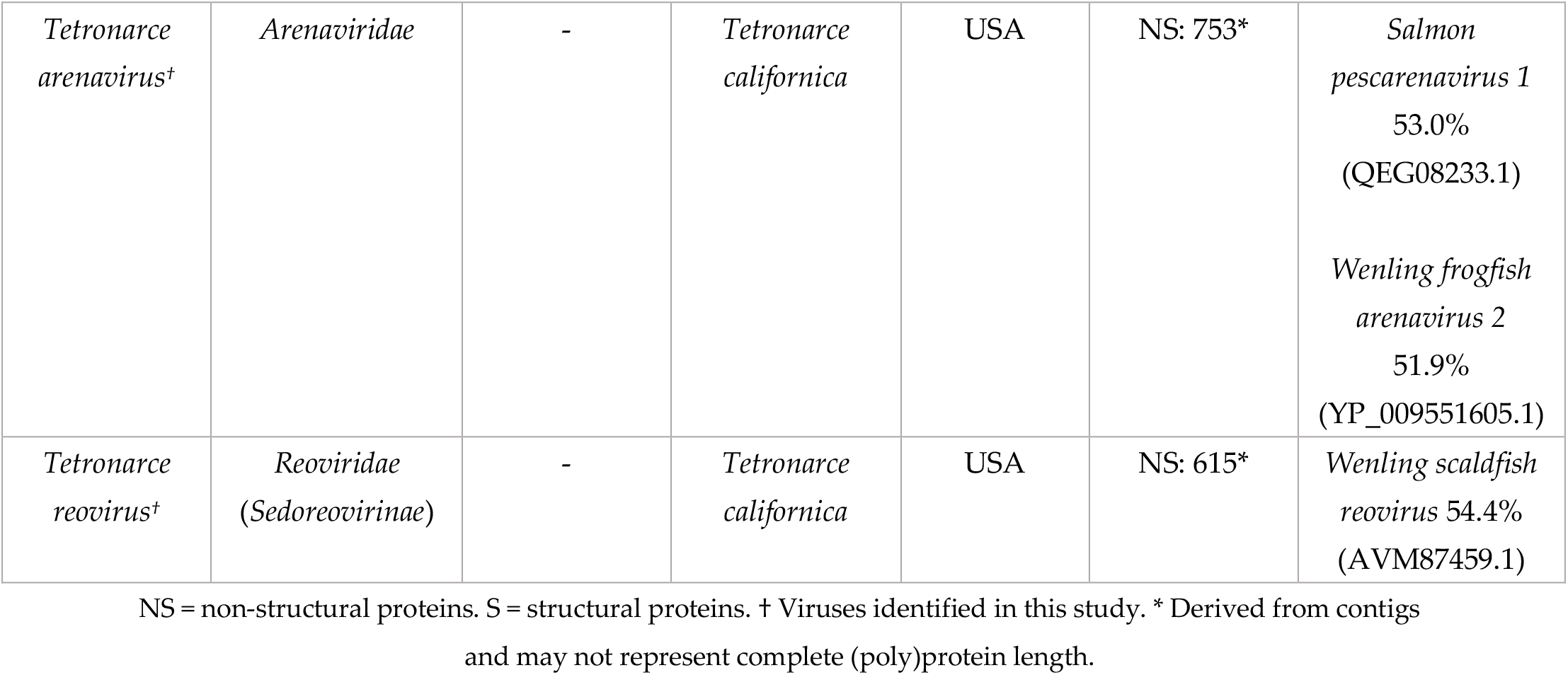
Novel Tetronarce viruses and *Matonaviridae* referenced in this study.

To recover further genomic regions of *Tetronarce matonavirus*, we *de novo* assembled the raw sequencing reads from the Pacific electric ray sample. Accordingly, transcripts of a non-structural polyprotein, the p90 (RdRp) protein (815 amino acids in length in RuV), were recovered. By mapping contigs to the ruhugu virus non-structural polyprotein reference sequence, a continuous 399 amino acid section of the RdRp was identified and used for additional phylogenetic analysis. The RdRp fragment matched most closely to the RdRp of RuV (accession: YP_004617078.1, identity: 64.9%, query coverage: 99%, e-value: 0.0, see Table 1).

### 3.2 Identification of novel Arenaviridae and Reoviridae in Pacific electric ray

To confirm that the host sequence reads analysed were indeed from the Pacific electric ray, we mined for additional viruses in the relevant transcriptome data. Accordingly, BLASTx analyses revealed contigs containing divergent sequences most closely related to viruses within *Arenaviridae* and *Reoviridae*. Specifically, we obtained a 753 amino acid fragment of the L segment of a novel arenavirus (tentatively termed *Tetronarce arenavirus*) and a 615 amino acid segment of the segment 1 protein of a novel reovirus (*Tetronarce reovirus*). BLASTx searches indicated that the 753 amino acid *Tetronarce arenavirus* fragment was most closely related to *salmon pescarenavirus 1* (53.0% sequence identity, Table 1) and the 615 *Tetronarce reovirus* segment was most closely related to *Wenling scaldfish reovirus* (54.4% sequence identity, Table 1).

### 3.3 Screening for host genes

To further address whether the host reads analysed were from the Pacific electric ray, rather than inadvertent mammalian contamination, the assembled Pacific electric ray contigs were screened for non-electric ray vertebrate annotations. We found that the closest genetic matches to all contigs with sequence homology to eukaryote genomes were either from the Pacific electric ray, related species within the Batoidea superorder, or genes that are highly conserved across vertebra, including cartilaginous and bony fish. Hence, these results strongly suggest that the *Tetronarce matonavirus* was indeed present in the Pacific electric ray transcriptome.

### 3.4 Virus abundance

We estimated the standardised abundances of the three novel viruses in comparison to the stably expressed host gene, 40S ribosomal protein S13 (RPS13) (Figure 1a). Of the three viruses present, *Tetronarce matonavirus* was the most abundant (standardised abundance = 4.4.×10^−6^), followed by *Tetronarce arenavirus* (8.0×10^−7^), and then *Tetronarce reovirus* (2.7×10^−7^). The abundance of reads from the host gene, RPS13, was 2.4×10^−4^.

**Figure 1.**
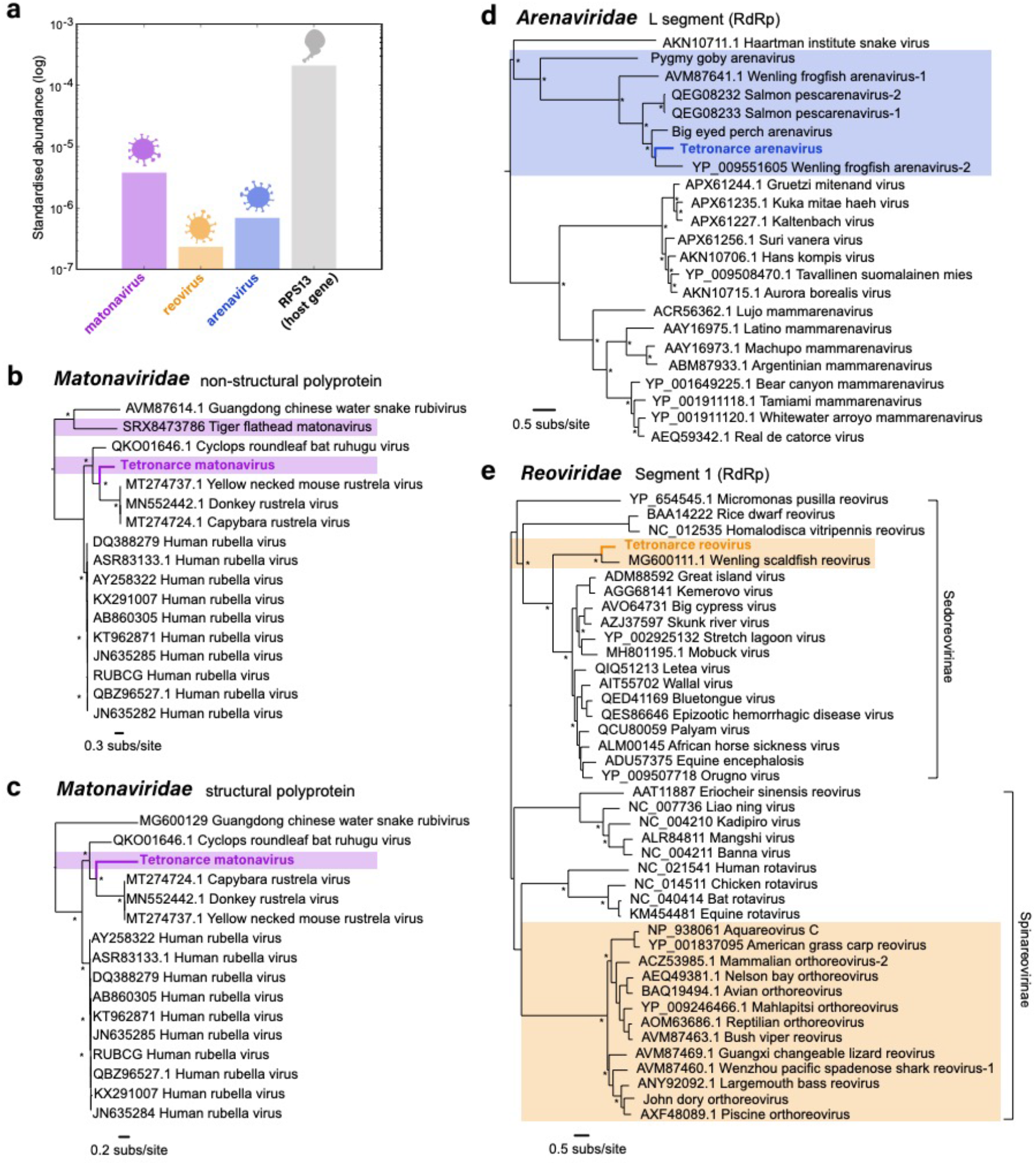
(a) Standardised abundance of viral reads (log) from the novel viruses (matonavirus, reovirus, and arenavirus) and a host gene (RPS13). **(b and c)** Phylogenetic trees of the non-structural and structural polyproteins of *Tetronarce matonavirus* within the *Matonaviridae*, respectively. **(d)** Phylogenetic tree of the L segment (RdRp) of *Tetronarce arenavirus* within the *Arenaviridae*. **(e)** Phylogenetic tree of segment 1 (RdRp) of *Tetronarce reovirus* within the *Reoviridae*. All phylogenetic trees are rooted at the midpoint. Viruses from fish species are highlighted within each phylogeny. Names of novel viruses identified in this study are in bold and highlighted. Nodes with bootstrap support values of >70% are indicated with asterisks (*).

### 3.5 Phylogenetic analysis of the novel viruses

Maximum likelihood phylogenetic trees were estimated using amino acid sequences of the non-structural (Figure 1b) and structural (Figure 1c) polyproteins of *Matonaviridae*, as well as the RdRp (L segment) of *Arenaviridae* (Figure 1d) and the RdRp (segment 1) of *Reoviridae* (Figure 1e) to determine the evolutionary relationships of the new Pacific electric ray viruses in the context of previously described virus species.

For phylogenetic analysis of *Tetronarce matonavirus*, separate sequence alignments were made for the non-structural and structural polyprotein and individual trees were generated for each genomic region. However, phylogenetic analysis of both the non-structural and structural polyproteins revealed similar topologies, with the novel *Tetronarce matonavirus* being most closely related to *Rustrela virus* (found in donkeys, mice, and capybara) and forming an ingroup to *Ruhugu virus* (found in bats). Strikingly, therefore, *Tetronarce matonavirus* does not group with the only other fish virus in this family, *Tiger flathead matonavirus*, and the evolutionary history of this virus family does not follow strict virus-host co-divergence.

In contrast, *Tetronarce arenavirus* falls within a group of viruses that have been identified in other fish host species, including *Big eyed perch arenavirus* and *Wenling frogfish arenavirus-2*. Unlike the *Reoviridae* and *Matonaviridae* phylogenetic trees, all fish arenaviruses identified to date form a distinct fish arenavirus clade.

The *Reoviridae* comprises two sub-families (*Sedoreovirinae* and *Spinareovirinae*). Reoviruses have previously been identified in fish (Figure 1e); however, these viruses are primarily Orthoreoviruses and Aquareoviruses (sub-family: *Spinareovirinae*). The *Tetronarce reovirus* and its closest match in the NCBI database (*Wenling scaldfish virus*, Table 1) are the only two fish viruses that fall within the *Sedoreovirinae* sub-family and appear to be more closely related to vector-borne reoviruses (orbiviruses) known to infect various mammalian hosts [28], including *African horse sickness virus* (AHSV) and *Equine encephalosis virus* (EEV), and the more recently discovered snake-associated *Letea virus* [29].

### 3.6 Matonaviridae amino acid conservation

We screened for evidence of conserved sequences or motifs between the novel matonavirus and previously described viruses. An immune-reactive region within the E1 structural protein of RuV [30] containing four neutralising B cell epitopes N1 – N4 [31] revealed significant levels of conservation with corresponding regions in the other mammalian matonaviruses [12]. These regions were also well conserved in *Tetronarce matonavirus* (Figure 2), compatible with our phylogenetic results. In contrast, the *Guangdong Chinese water snake rubivirus*, which has a very short structural protein sequence (687 amino acids compared to ∼1098 amino acids), exhibited little conservation of the E1 protein and other structural proteins overall. There is clear conservation in regions corresponding to epitopes N2 – N4 between the mammalian viruses and the electric ray virus, including a glycine residue in N4 conserved across all five viruses in the alignment (Figure 2). In addition, a highly conserved glycine-aspartic acid-aspartic acid (GDD) motif present in the RdRp [32] was also present and conserved in *Tetronarce matonavirus* as well as all other matonaviruses referenced here (Figure 2).

**Figure 2.**
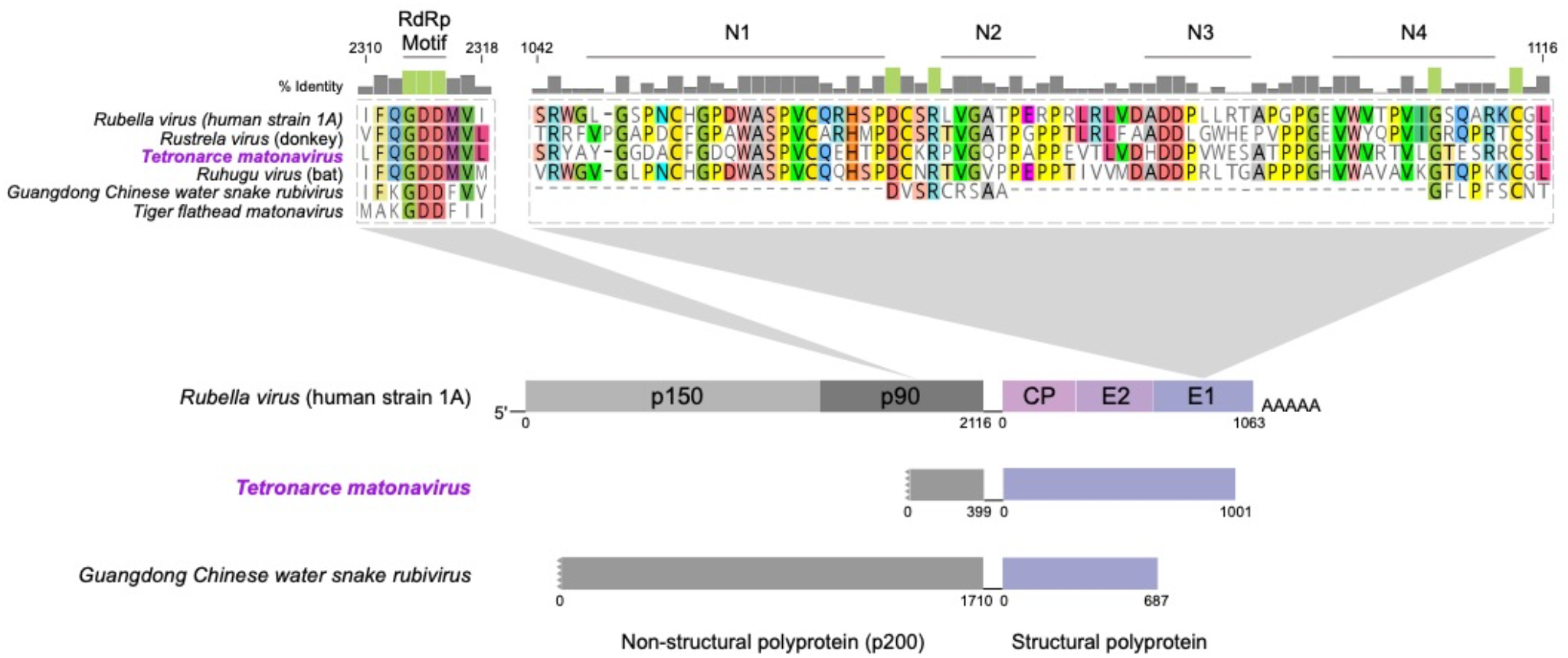
Conserved sequence motifs between human *Rubella virus* (strain 1A), *Rustrela virus* (donkey), *Ruhugu virus* (bat), *Tetronarce matonavirus, Tiger flathead matonavirus*, and *Guangdong Chinese water snake rubivirus* shown. The non-structural (p200) polyprotein contains a conserved GDD amino acid motif in the p90 (RdRp) protein at positions 2313 – 2315. Regions corresponding to RuV B cell epitopes N1 – N4 within E1 in the structural protein alignment at positions 1042 – 1116 are also illustrated. Bar graphs of percentage identity (%) of residues shown above the alignments with identities of 100% highlighted green. Genomic organisation of *Matonaviridae* (*Rubella virus* strain 1A) is shown below with segments able to be recovered from *Tetronarce matonavirus* and *Guangdong Chinese water snake rubivirus* indicated underneath for comparison. These comprise the complete structural polyprotein and partial non-structural polyproteins.

## 4. Discussion

We identified a complete structural polyprotein and a partial non-structural polyprotein (comprising a section of the RdRp) from a divergent matonavirus in a transcriptome of a Pacific electric ray. Additionally, partial RdRp transcripts of other novel viruses within the *Reoviridae* and *Arenaviridae* were identified in this same fish transcriptome.

The novel *Tetronarce matonavirus* identified here exhibited clear sequence similarity to RuV, as well as to the *Rustrela* and *Ruhugu viruses* recently discovered in other mammals. High levels of sequence conservation within the E1 structural protein have been previously shown among RuV, *Rustrela virus*, and *Ruhugu virus* [12], including regions corresponding to various immune-reactive B and T cell epitopes in RuV [30, 31]. *Tetronarce matonavirus* shares comparable levels of amino acid conservation to mammalian matonaviruses in this region of E1, whereas the other matonavirus from a non-mammalian host (*Guangdong Chinese water snake virus*) does not. However, we were unable to recover a structural polyprotein from *Tiger flathead matonavirus* to incorporate it into our analysis. The non-structural fragment of the novel virus also contains a highly conserved GDD motif within the RdRp, which is highly conserved across many RNA viruses and is essential for efficient replication [32].

Of particular note was that phylogenetic analysis suggested that *Tetronarce matonavirus* is more closely related to *Rustrela virus* than to the other, more divergent, fish matonavirus, and hence is indicative of a complex history of cross-species transmission events. Such a conclusion is not dependent on the position of the tree root. Given that this phylogenetic position is somewhat paradoxical, it is important to exclude sample contamination, or that the *Tetronarce matonavirus* was in fact obtained from a component of the diet of *T. californica*. Importantly, no significant non-fish matches were identified in the *T. californica* transcriptome library, with most host reads producing matches to *T. californica* or related fish species. Similarly, that the novel arenavirus and reovirus identified were most closely related to previously documented fish-associated viruses, rather than those from mammals, again suggests that the unusual phylogenetic position of *Tetronarce matonavirus* is *bona fide*. Hence, these data are indicative of a complex history of cross-species transmission of viruses in the *Matonaviridae*, potentially involving cross-species transmission events among different vertebrate classes. Evidence of such expansive host-jumping events has been observed in the evolution of other virus families, such as hepatitis B viruses in the *Hepadnaviridae*, in which a number of fish viruses group closely with those identified in mammals [33]. Indeed, cross-species transmission is commonplace in RNA viruses as a whole [34], and the expansion of the *Matonaviridae* with viruses from such a broad host range suggests this process has also played an important role in the evolutionary history of these viruses.

A potentially new reovirus -*Tetronarce reovirus*, was also identified in our analysis. The majority of reoviruses found in fish fall within the sub-family S*pinareovirinae*. In contrast, *Tetronarce reovirus* appears to be one of two viruses currently known that fall within the S*edoreovirinae* sub-family that include vector-borne orbiviruses such as AHSV and EEV that cause fatal diseases in equines [35, 36]. However, the host range of this sub-family is continually expanding and now contains a recently identified snake-associated *Letea virus* [29]. It is also notable that *Tetronarce reovirus* is most clearly related to another fish-associated reovirus -*Wenling scaldfish reovirus*. Similarly, the *Arenaviridae* has also been recently expanded, with the host range thought to be previously limited to rodents and humans, now includes other warm-and cold-blooded vertebrates [37]. The novel arenavirus identified here -*Tetronarce arenavirus* – clusters with a group of fish and amphibian viruses.

The recovery of a complete structural polyprotein and a partial RdRp, both with conserved amino acid sequences and motifs present across *Matonaviridae*, as well as a relatively high viral abundance compared with a stably expressed host gene, is highly suggestive of the existence a novel rubi-like virus in *T. californica*. While screening of annotated contigs and the existence of two other potentially novel viruses sharing sequence identity to other fish viruses in the assembled transcriptome suggest the Pacific electric ray to be the true host of these viruses, further surveying of this species for viral genomes would be beneficial to further understanding the origin and emergence of matonviruses like RuV. Overall, these findings expand the currently established host range of the *Matonaviridae* and suggest a more complex evolutionary history than previously suspected. More broadly, this study demonstrates the value of performing extensive transcriptome sequencing and screening of a wide range of potential hosts to identify animal relatives of viruses of significant health concern and to characterise more of the global virosphere.

## Author Contributions

Conceptualization, R.M.G., E.C.H. and J.L.G.; formal analysis, R.M.G., E.C.H. and J.L.G.; resources, R.M.G., E.C.H. and J.L.G.; writing—original draft preparation, R.M.G., E.C.H. and J.L.G.; writing— review and editing, R.M.G., E.C.H. and J.L.G.; funding acquisition, E.C.H. and J.L.G. All authors have read and agreed to the published version of the manuscript.

